# The C3-C3aR axis modulates trained immunity in alveolar macrophages

**DOI:** 10.1101/2024.11.01.621042

**Authors:** Alexander P. Earhart, Alberto Lopez, Josue I. Hernandez, Aasritha Nallapu, Deebly Chavez, Sayahi Suthakaran, Brian Yang, Jungheun Hyun, Rafael Aponte Alburquerque, Marick Starick, Lorena Garnica, Ayse Naz Ozanturk, Rahul Kumar Maurya, Xiaobo Wu, Jeffrey A. Haspel, Jae Woo Lee, Jaime Hook, Hrishikesh S. Kulkarni

**Affiliations:** Divisions of Pulmonary and Critical Care Medicine, Icahn School of Medicine at Mount Sinai, New York, NY, USA; Divisions of Rheumatology, John T. Milliken Department of Medicine, Washington University School of Medicine, Icahn School of Medicine at Mount Sinai, New York, NY, USA; Departments of Medicine, Icahn School of Medicine at Mount Sinai, New York, NY, USA; Departments of Anesthesia, University of California, Los Angeles David Geffen School of Medicine, Icahn School of Medicine at Mount Sinai, New York, NY, USA; Lung Imaging Laboratory, Division of Pulmonary, Critical Care, and Sleep Medicine, Department of Medicine, Icahn School of Medicine at Mount Sinai, Icahn School of Medicine at Mount Sinai, New York, NY, USA; Department of Microbiology, Icahn School of Medicine at Mount Sinai, New York, NY, USA; Department of Stem Cell Biology and Regenerative Medicine, Icahn School of Medicine at Mount Sinai, New York, NY, USA; Global Health and Emerging Pathogens Institute, Icahn School of Medicine at Mount Sinai, New York, NY, USA

**Author notes:** co-first author (Alexander P. Earhart, Alberto Lopez). **Corresponding author(s):** Alexander P. Earhart, PhD, Washington University School of Medicine, Hrishikesh S. Kulkarni, MD, MSCI, UCLA David Geffen School of Medicine. **Contributions:** A.P.E. was involved in conceptual development, methodology, investigation, statistical analysis, writing – original draft, writing – review and editing; and funding acquisition; A.E.L. and A.N. were involved in methodology, investigation, statistical analysis, writing – original draft, writing – review and editing; D.C., S.S., B.Y., R.A., M.S., A.N., L.G., A.N.O., R.K.M., X.W., J.A.H., and J.H. were involved in the methodology, investigation, and writing – review and editing; H.S.K. was involved in conceptual development, methodology, investigation, statistical analysis, writing – original draft, writing – review and editing; supervision, and funding acquisition.

## Abstract

Complement protein C3 is crucial for immune responses in mucosal sites such as the lung, where it aids in microbe elimination and enhances inflammation. While trained immunity – enhanced secondary responses of innate immune cells after prior exposure – is well-studied, the role of the complement system in trained immune responses remains unclear. We investigated the role of C3 in trained immunity and found that alveolar macrophage *C3* and *C3aR1* expression increased in humans after an intranasal exposure to a training stimulus. *In vivo*, trained wild-type mice showed significantly elevated pro-inflammatory cytokines and increased C3a levels upon a second stimulus. *Ex vivo*, trained C3-deficient alveolar macrophages (AMs) displayed reduced chemokine and cytokine output as well as impaired phagocytosis and reactive oxygen species (ROS) production compared to wildtype AMs. Real-time confocal microscopy of live, intact mouse alveoli revealed that AMs internalize C3 rapidly after alveolar microinstillation, as compared to C3a. Correspondingly, the blunted cytokine output was restored by exogenous C3 but not by C3a. Inhibiting C3aR, both pharmacologically and with a genetic C3aR knockout, prevented this restoration, indicating the necessity of C3aR engagement. Mechanistically, trained WT AMs demonstrated enhanced glycolytic activity compared to C3-deficient AMs – a defect corrected by exogenous C3 in a C3aR-dependent manner. These findings reveal that C3 modulates trained immunity in AMs through C3aR signaling and highlight a novel role for C3 in trained immunity.

## INTRODUCTION

The complement system is a crucial part of immunity, consisting of proteolytic enzymes that generate fragments which enhance antibody binding and help induce phagocytosis (Sahu et al., 2022). A central component of this system is the C3 protein, which is cleaved into its constituent components C3a and C3b upon activation (Kulkarni et al., 2018). C3b tags pathogens for phagocytosis and plays a vital role in the formation of the C3 convertase, which cleaves C5 and eventually forms the membrane attack complex that lyses targeted cells (Janssen et al., 2006). Meanwhile, C3a acts as an anaphylatoxin, binding to its cognate C3a receptor (C3aR) on and within cells, and modulates inflammatory responses (Kildsgaard et al., 2000). C3 activation has also been shown to favor an increase in glycolysis, suggesting a plausible mechanism for enhanced inflammatory activity (Friščić et al., 2021; S. Kang et al., 2024). Moreover, C3 is present at low levels in the bronchoalveolar lavage (BAL) fluid of uninjured mice as well as humans, indicative of localized production (Bolger et al., 2007). This local C3 production increases during injury, affecting mucosal responses to infection (Bolger et al., 2007). Additionally, the conversion of C3 to a conformationally altered C3(H_2_O) moiety increases during inflammation (Elvington et al., 2019). This C3(H_2_O) form is that which is internalized by various cell types, playing a key role in modulating survival and effector immune responses (Elvington et al., 2017; Kulkarni et al., 2019). Thus, local C3 activity plays an important role in modulating mucosal immune responses (Kulkarni et al., 2024).

“Trained immunity” refers to cells such as innate immune cells (i.e., monocytes and macrophages) exhibiting robust, enhanced inflammatory responses upon secondary stimulation after a prior insult (Quintin et al., 2012; Saeed et al., 2014). This form of “memory” is considered broadly antigen nonspecific, yet lasts several months after the initial stimulus (Netea et al., 2020). Underlying the induction of trained immunity are epigenetic and metabolic reconfigurations occurring after an initial stimulus, which prime transcriptional machinery for rapid activation after engaging secondary insults (Bekkering et al., 2014; Fanucchi et al., 2021). For instance, the bacterial and fungal cell wall component 1,3-D-β-glucan, binds the dectin-1 pattern recognition receptor, which induces metabolic changes favoring persistent glycolytic activity (Cheng et al., 2014; Earhart et al., 2023). This, in turn, promotes shifts in epigenetic modifications such as histone acetylation and methylation favoring strong pro-inflammatory gene expression upon restimulation, and can be long-lasting (Fok et al., 2019; Tercan et al., 2021). While knowledge of trained immunity is primarily systemic, there is growing evidence for site-specific effects such as in alveolar macrophages (AMs) (Chakraborty et al., 2023; Zahalka et al., 2022). Little is currently known about how the complement system affects trained immunity. Here, we sought to investigate whether the C3 protein – a key component of immune responses at mucosal sites such as the respiratory system – affects trained immunity in AMs and determine whether any effect is mediated by C3aR activity.

## RESULTS

To assess the relevance of C3 in response to a training stimulus, we interrogated a publicly available dataset of bronchoalveolar lavage specimens from human volunteers who underwent aerosolized Bacillus Calmette Guerin (BCG) administration (Marshall et al., 2025). Both the mean expression as well as the percentage of AMs expressing *C3* and *C3aR1* increased at day 2 post-BCG exposure, and remained elevated at day 7, as compared to their expression in AMs of volunteers who received saline (Fig. 1A-C). *C3* expression was highest in the activated alveolar macrophage subset (Supplementary Fig. S1A-C). To investigate the role of C3 in pulmonary immune cell-trained immunity, we inoculated C57BL/6J wild-type (WT) and B6.129S4-*C3*^*tm1Crr*^/J C3 knockout (C3KO) mice intranasally with heat-killed *Pseudomonas aeruginosa* (HKPA) for training or vehicle control (PBS, untrained). After 14 days, lipopolysaccharide (LPS) from *Escherichia coli* was also administered intranasally to both mouse strains to induce secondary stimulation for 24 h, followed by euthanasia and bronchoalveolar lavage (BAL) as previously described (Fig. 1D) (Sahu et al., 2023). BAL proinflammatory chemokines and cytokines (CXCL1, CXCL2, IL-6, and TNFα) were quantified using ELISAs. Additionally, C3a, which is generated when C3 is activated and cleaved, was also measured using an ELISA specific to its neo-epitope. CXCL1, CXCL2, IL-6, and TNFα were all significantly elevated in trained versus untrained WT BAL (Fig. 1E). Levels of C3a were increased in trained versus untrained WT BAL, indicating enhanced C3 activation is part of the trained immune response (Fig. 1F). The enhancement in BAL IL-6 and TNFα with training was blunted in C3-deficient mice compared to WT mice (Fig. 1G). Of note, no difference was observed in BAL protein, neutrophils and cytokines (i.e., CXCL1, TNFα) at day 14 (prior to LPS exposure) in C3-deficient compared to wild-type mice, regardless of prior HKPA (training) exposure (Fig.S1D-G), suggesting the resolution of acute inflammation. These results suggest C3 is required for trained immunity *in vivo*, and that the observed responses to the secondary challenge are not confounded by persistent inflammation from the initial exposure.

**Figure 1.**
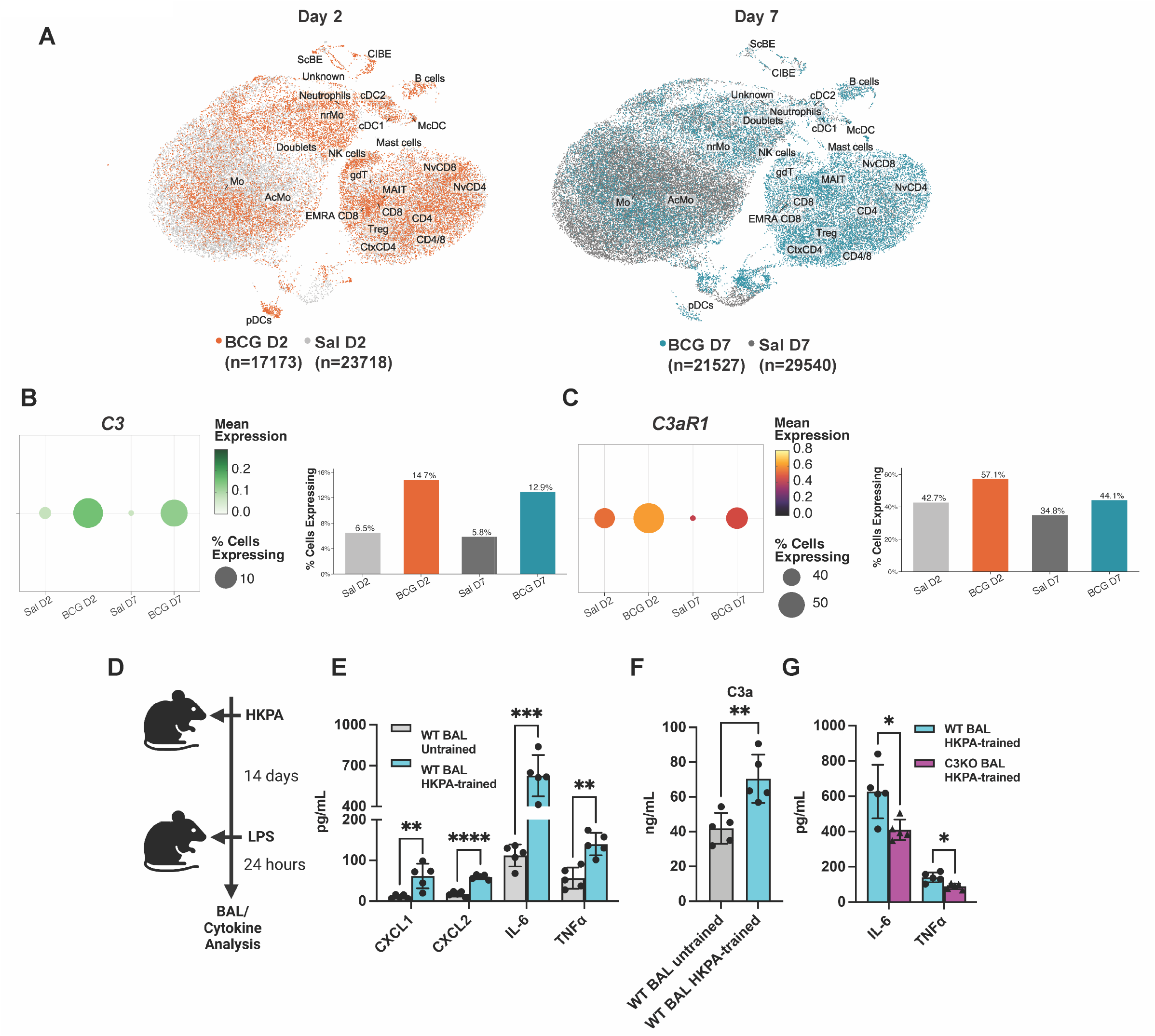
C3 deficiency predisposes to impaired pulmonary trained immunity. (A) UMAP (Uniform Manifold Approximation and Projection) plots showing the identity of each cell cluster in human bronchoalveolar lavage (BAL) from BCG- and saline (Sal)-treated donors at Day 2 (BCG n=17,173; Sal n=23,71S) and Day 7 (BCG n=21,527; Sal n=29,540) post-vaccination. N=3 volunteers in each group. Cell clusters include: macrophage (Mo), activated macrophage (AcMo), non-resident macrophage (nrMo), **NK** cells, γ δ T cells, plasmacytoid DC (pDC), conventional DC (cDC1, cDC2), migratory DC (McDC), B cells, MAIT cells, CD4, CDS, CD4/S, cytotoxic-like CDS T cells (CtxCD4), terminally differentiated effector memory CD45RA-re-expressing CDS T cells (EMRA CDS), naÏfve T cells (NvCD4, NvCDS), T regulatory cells (Treg), neutrophils, mast cells, doublets, ciliated bronchial epithelial cells (GIBE), secretory bronchial epithelial cells (ScBE), and unidentified cells (Unknown). Data from Marshall et al, 2025. (B) Dot plot and bar chart showing C3 expression across all human alveolar macrophages (Mcp; Mo, AcMo, nrMo pooled) across conditions. Dot size reflects the percentage of cells expressing C3; dot color reflects mean normalized expression. Bar chart shows percentage of C3-expressing M φ per condition. (C) As in (B), for C3AR1. BCG vaccination increases the proportion of C3AR1-expressing alveolar macrophages at both timepoints relative to saline controls. (D) Schematic representing the training of mice via the intranasal route with heat-killed Pseudomonas aeruginosa (HKPA) and subsequent restimulation with lipopolysaccharide (LPS), followed by BAL and cytokine analysis. Created with BioRender. (E) *WT* untrained compared against WT-trained BAL levels of CXCL1, CXCL2, IL-6, and TNF α. Comparison of BAL C3a levels, similar to (E). (G) WT-trained versus C3-deficient (C3KO)-trained BAL concentrations of IL-6 and TNFα. WT-trained levels derived from (E) for comparison with C3KO-trained mice. Data were compared with two-sided unpaired t-tests with (E,G) or without (F) Holm-Sidak correction for multiple hypothesis testing. Each point represents a measurement from one mouse with at least n=4 in each group, mean± SD shown. *p < 0.05, **p < 0.01, ***p < 0.001.

To examine how C3 affects trained immunity specifically in tissue-resident phagocytes, we set up an *ex vivo* culture system using primary alveolar macrophages (AM) from C3-deficient and wildtype mice (Gorki et al., 2022). Like the *in vivo* experiments (Fig. 1), we used HKPA to induce trained immunity in cultured AMs, followed days later by LPS challenge and analysis of cytokine secretion in the presence or absence of C3 deficiency (Fig. 2A). Supporting our *in vivo* results, training by HKPA was sufficient to augment LPS-induced proinflammatory cytokine production from primary AMs (Fig. 2B). However, C3-deficient AMs demonstrated blunted LPS-induced proinflammatory cytokine production post-training, as measured by IL-6 and TNFα secretion (Fig. 2C) as well as impaired phagocytosis and reactive oxygen species (ROS) production (Fig. S2A-C). Employing heat-killed *Candida albicans* (HKCA) in place of HKPA produced similar results (Fig. 2D-G). To assess if the in vivo training specifically influenced AMs, we also trained the mice in vivo using HKPA (using PBS as a control for 14 days, then harvested AM, and provided the second stimulus (LPS) ex vivo. In vivo trained AM from C3-deficient demonstrated a blunted cytokine response to LPS ex vivo compared to AM from wildtype mice (Fig. S2D), These data suggest a specific role for C3 in AM immune training, consistent with our *in vivo* observations.

**Figure 2:**
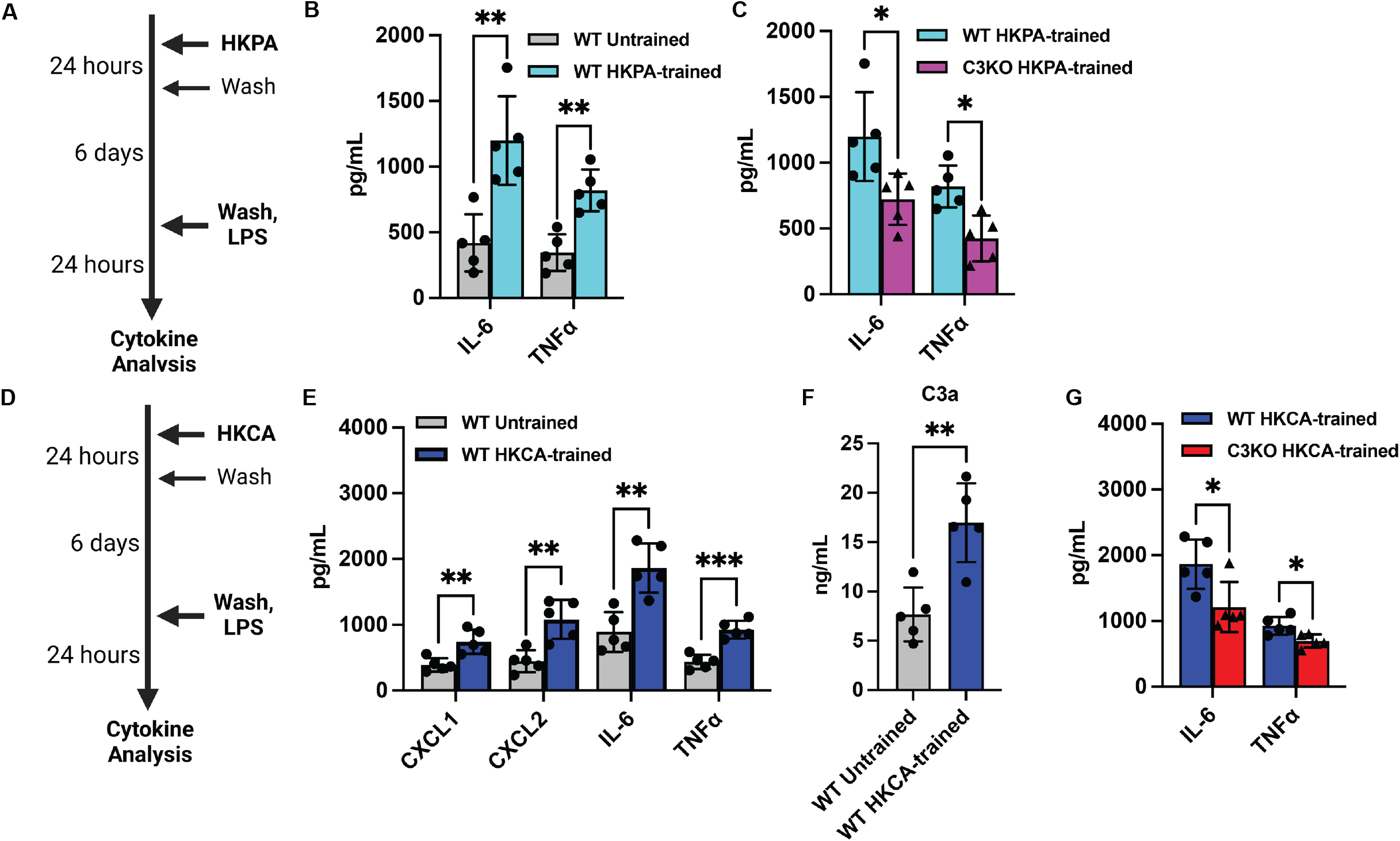
C3 deficiency results in impaired trained immune responses in ex vivo alveolar macrophages (AMs). (A) Schematic representing in vitro training of AMs with HKPA, with later stimulation by LPS and subsequent cytokine analysis of the supernatants. Created with BioRender. (B-C) Effects of HKPA-induced training in vitro on IL-6 and TNFα in supernatant from (B) WT AM, and (C) their comparison with C3KO-trained AMs. (D) Schematic representing in vitro training of AMs with HKCA, with subsequent restimulation by LPS and cytokines in supernatant. Created with BioRender. (E) Effects of heat-killed Candida albicans (HKCA)-induced training in vitro on cytokines in supernatant from WT AM. (F) Comparison of C3a levels post-HKCA training, similar to (B). (G) Comparison of IL-6 and TNFα post-HKCA training in WT versus C3KO AMs. WT-trained levels derived from (D) for comparison with C3KO-trained AMs. Data were compared with two-sided unpaired t-tests with (B,C,E,G) or without (F) Holm-Šidák correction for multiple hypothesis testing. Each point is a technical replicate from pooling AMs from at least n=4 mice in each group, with mean ± SD shown. Each experiment repeated twice. *p < 0.05, **p < 0.01, ***p < 0.001.

In addition to synthesizing C3, multiple cell types internalize C3 as C3(H_2_O) in vitro, which promotes cell survival and effector responses (Elvington et al., 2017; Kulkarni et al., 2019). To assess if C3 uptake by AMs can overcome the defect in responses from C3-deficient AMs, we first interrogated if mouse AMs internalize C3 in vivo. Labeled C3a was separately used as a control. Real-time confocal microscopy of live, intact mouse alveoli (Fig. 3A&B) revealed that CD11c^+^ AMs internalize fluorescently labeled C3 rapidly after alveolar microinstillation, as compared to C3a (Fig.3C-G). Incubation of human precision-cut lung slices with labeled C3 revealed that human AM also internalize C3 in situ, thereby, validating our observations from the intact mouse lung using human lung tissues (Fig. S3).

**Figure 3.**
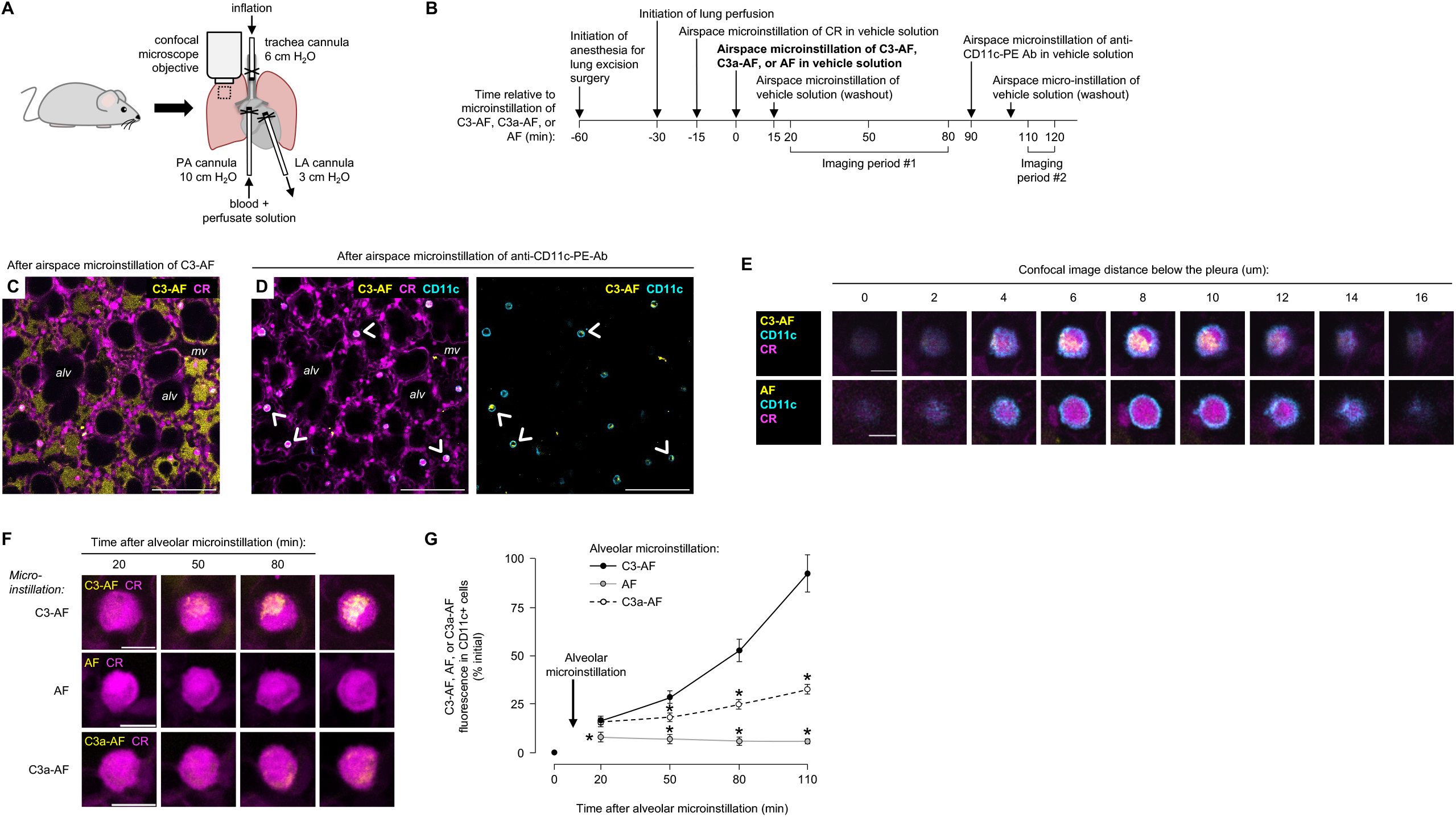
Alveolar macrophages take up C3 from airspaces of live alveoli in situ. Cartoons in A-B show the experimental design of the confocal imaging studies shown in C-G. As indicated in A-B, we microinstilled alveolar airspaces of live, intact, perfused mouse lungs sequentially with: cell-permeant calcein red-orange dye (*CR*); Alexa Fluor 647 (*AF*), AF-tagged C3 (*C3-AF*), or AF-tagged C3a (*C3a-AF*); and phycoerythrin *(PE*)-tagged anti-CD11c Ab. The confocal image in C shows C3-AF fluorescence (*yellow*) in alveolar airspaces and CR fluorescence (*magenta*) in airspace-facing cells, including the alveolar epithelium and alveolar macrophages. *alv*, example airspace; *mv*, microvessel. Confocal images in D show the same alveoli, but CD11c fluorescence (*cyan*) now marks CD11c+ cells. *Arrowheads* point out example CD11c+ cells with intracellular C3-AF fluorescence. High power confocal images of CD11c+ cells (E-F) and group data (G) show C3-AF accumulated in cytosols of CD11c+ cells over time. In G, *circles* indicate mean ± SEM fluorescence in all of the CD11c+ cells present in imaging fields of at least 30 alveoli; *n* = 4 microinstillations in 2 lungs per group; \**p* < 0.05 versus C3-AF by ANOVA with post hoc Tukey testing. C3-AF, C3a-AF, and AF fluorescence in airspaces was normalized to C3-AF, C3a-AF, and AF fluorescence in glass micropipettes. Scale bars: 100 (C-D) and 10 (E-F) µm.

To mechanistically validate the role of C3 uptake in the training of AMs, we cultured C3-deficient AMs with exogenous C3 at a dose that leads to the cellular uptake of C3(H_2_O) (15 µg/mL, Fig 4A) (Elvington et al., 2017; Kulkarni et al., 2019). As a control, we incubated cells with labeled C3a, which is not internalized rapidly (Mogilenko et al., 2022). As measured by IL-6 and TNFα after a secondary stimulus, exogenous C3 protein restored the trained responses by AM to WT levels (Fig. 4B). In contrast, these responses were not restored to WT levels by exogenous C3a, which stays outside the cell (Fig. 4C).

**Figure 4.**
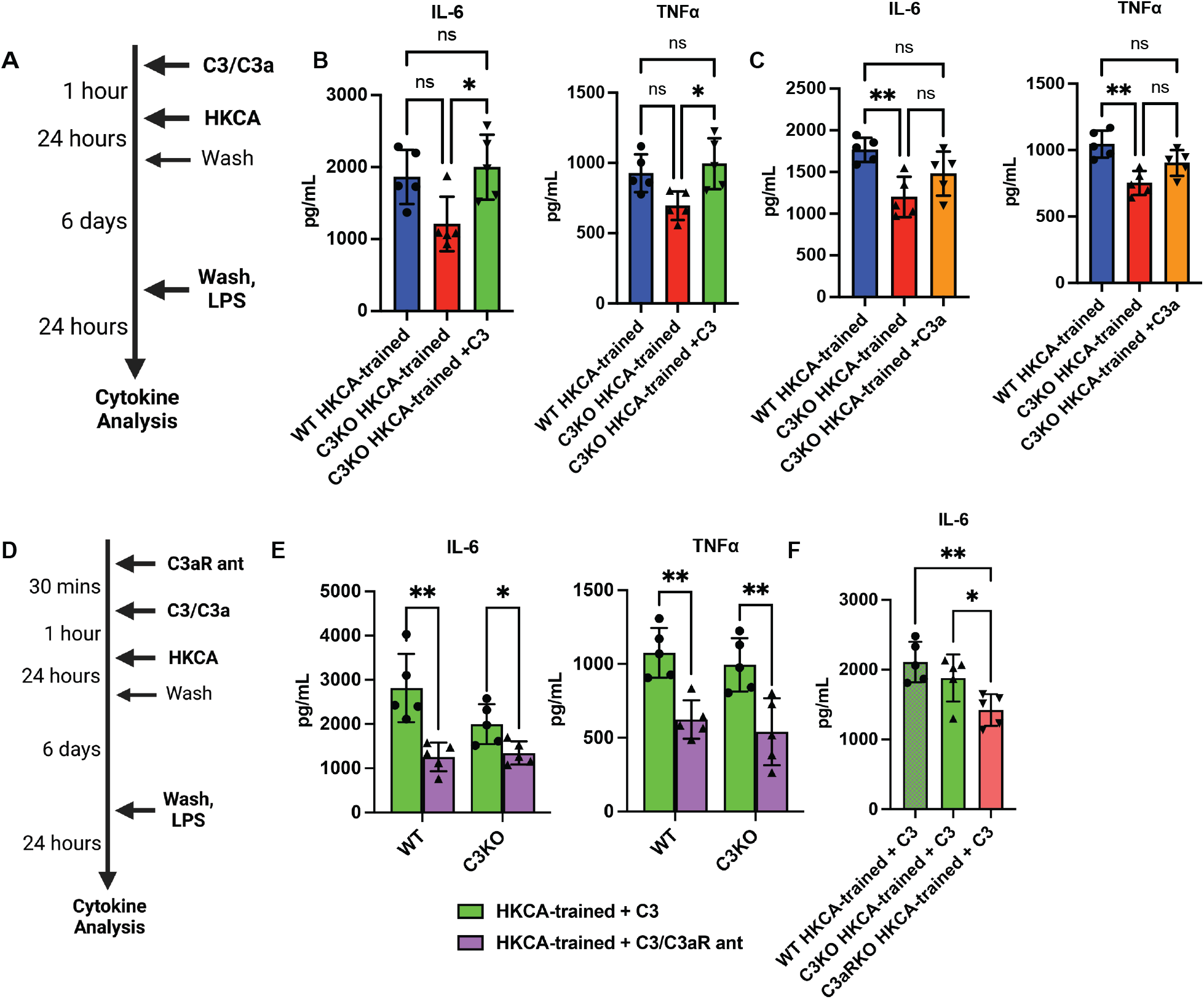
C3 uptake enhances trained immune responses in ex vivo alveolar macrophages (AMs) via the C3a receptor (C3aR). (A) Schematic representing in vitro training of AMs with HKCA, with pre-treatment of C3 or C3a prior to induction of training, and later stimulation by LPS and subsequent cytokine analysis of the supernatants. Created with BioRender. (B) Effects of adding C3 prior to training on IL-6 and TNFα levels from C3KO AMs and their comparison with WT-trained AMs. (C)Effects of adding C3a prior to training, similar to (B).(D) Schematic representing addition of the C3aR antagonist prior to C3 treatment and in vitro training of AMs with HKCA, with later stimulation by LPS and subsequent cytokine analysis of the supernatants. Created with BioRender. (E) Effects of C3aR antagonism on IL-6 and TNFα levels from trained WT and C3KO AMs treated with exogenous C3. (F) Comparison of IL-6 levels post-HK-CA-training in C3aR-deficient (C3aRKO), C3KO and WT AMs treated with exogenous C3. Data were compared using one way ANOVA with Dunnett’s post hoc tests (B,C,F) or two-sided unpaired t-testing with Holm-Šidák correction for multiple testing (E). Each point is a technical replicate made by pooling AMs from at least n=4 mice in each group, with mean ± SD shown, and each experiment was repeated twice. *p < 0.05, **p < 0.01.

Upon internalization, C3 is cleaved to C3a (Elvington et al., 2017). Intracellular C3a binds to C3aR and polarizes cytokine production in CD4^+^ T cells (Liszewski et al., 2013). To investigate whether a similar pathway could be important to C3-mediated trained immunity, we pre-treated AMs with a cell-permeable C3aR antagonist (SB290157) before immune training (Fig. 4D). C3aR antagonism blunted LPS-induced cytokine production in both WT and C3-deficient AMs that had previously been rescued with exogenous C3 (Fig. 4E). Importantly, genetic C3aR deficiency phenocopied the blunting of cytokine production seen in C3-deficient cells. Specifically, this defect in cytokine production post-training could not be rescued by exogenous C3 in C3aR-deficient AMs (Fig. 4F). Altogether, these findings support a causal role for C3 in AM trained immunity via the C3aR.

To explore transcriptomic changes relevant to how C3 influences trained immune responses, we performed bulk RNASeq comparing trained WT to trained C3KO AMs. C3-deficiency led to differential expression of not only innate immune genes but also several genes involved in metabolism post-training, including those relevant to glycolysis such as *gnpda1* and *aldoc* (Fig. 5A, Table S1). These metabolism-linked genes are significant because prior studies show that enhanced glycolytic metabolism is critical for trained immunity (Cheng et al., 2014). Therefore, we investigated whether the absence of C3 affects glycolytic flux in AMs (Fig. 5B). As measured by extracellular acidification rate, untrained C3-deficient AMs had comparable basal and maximum glycolytic flux compared to WT but failed to augment glycolysis upon training with HKCA (Fig. 5C). We next tested the ability of exogenous C3 to rescue the induction of training-associated glycolysis in C3-deficient AMs (Fig. 5D). Exogenous C3 returned glycolysis induction to the level of trained WT cells, suggesting intracellular processing of C3 is important for this effect (Fig. 5E). Like cytokines (Fig. 4), inhibiting C3aR prevented exogenous C3 from rescuing glycolysis induction during immune training (Fig. 5E).

**Figure 5.**
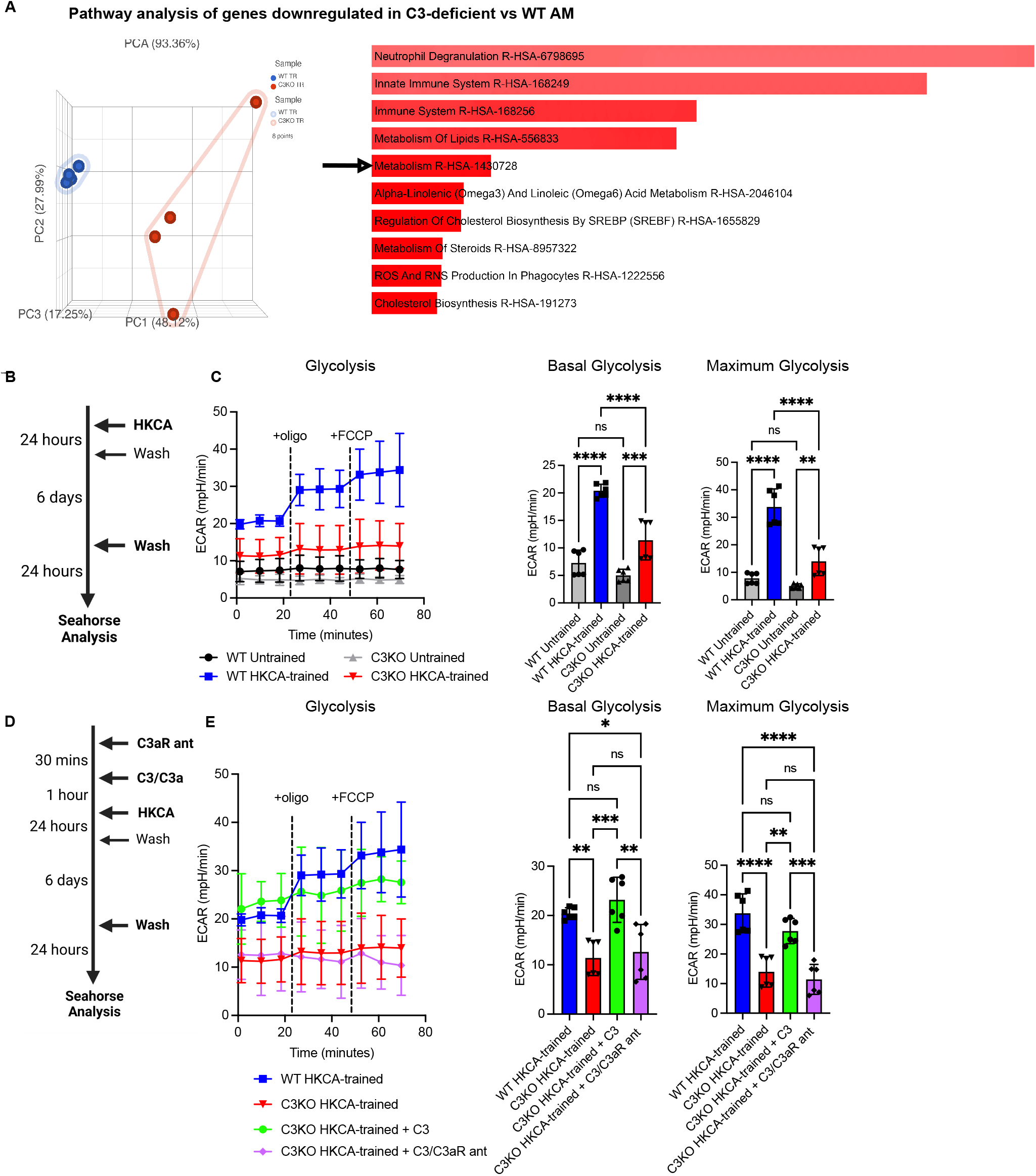
C3-C3aR axis is required for glycolysis as a part of trained immune responses in alveolar macrophages (AMs). (A) Principal component analysis (PCA, left) and EnrichR analysis of 391 genes (right, Table S1) downregulated in HKCA-trained C3KO vs WT AM by filtering genes (FDR step up ≤0.05). Arrow shows metabolism gene set in EnrichR; bars ranked by p-value. (B) Schematic representing in vitro training of AMs with HKCA. Created with BioRender. (C) Extracellular acidification rate (ECAR) from Seahorse analysis representing full glycolytic activity, and basal and maximum glycolysis in untrained and HKCA-trained WT and C3KO AMs. (D) Schematic representing addition of the C3aR antagonist (SB290157) prior to C3 treatment and in vitro training of AMs with HKCA. Created with BioRender. (E) Seahorse analysis in the presence and absence of exogenous C3 supplementation and C3aR antagonism. Each point is a technical replicate of pooled AMs from at least n=4 mice in each group, with mean ± SD shown. *p < 0.05, **p < 0.01, ***p < 0.001, ****p < 0.0001 when analyzed using one way ANOVA with Dunnett’s post hoc tests.

## DISCUSSION

Our data suggests C3 is upregulated after an initial inhalational exposure, and is required for effective reprogramming of AMs. In its absence, AMs demonstrate blunted immune and metabolic responses to a subsequent stimulus, which are rescued by exogenous C3 and are C3aR-dependent. Of note, an intracellular C3aR has been reported by several groups (Liszewski et al., 2013; Quell et al., 2017; Zha et al., 2019). We show that C3 is rapidly internalized by AMs, similar to other immune and non-immune cell types (Elvington et al., 2017; Ishii et al., 2021; Kulkarni et al., 2019), whereas C3a in itself is not internalized. We propose that in AMs, the internalization of C3(H_2_O) provides an intracellular source of C3a (first suggested in (Elvington et al., 2017)), which potentially engages with the C3aR to result in reprogramming. While our data suggests that exogenously added C3a does not significantly affect AM reprogramming in our model system, we cannot rule out that other C3-mediated paracrine signaling mechanisms may also be at play. Regardless, our data indicate that C3 makes a mechanistic contribution to trained immunity in AMs and implicate engagement of the C3aR as part of the pathway. We also acknowledge that other immune cells such as dendritic cells and neutrophils, and non-immune cells such as epithelial cells, endothelial cells and fibroblasts can also be involved in immune training (Bigot et al., 2025; Friščić et al., 2021; Moorlag et al., 2020).

Strengths of our study include *in vitro* and *in vivo* approaches to inducing trained immunity, orthogonal endpoints to quantify immune training (cytokine and ROS production, phagocytosis and metabolic readouts), and mechanistic gain- and loss-of-function experimental approaches. Future work involves identifying the downstream effectors of C3 and C3aR in the AM training process and their connection to immunometabolism. We also need to establish whether C3a-C3aR-mediated AM reprogramming can lead to altered disease outcomes. Systemically administered β-glucan has been shown to induce peripheral trained immunity and aggravate lung injury (Prevel et al., 2025) similar to disease in models of periodontitis and arthritis (Haacke et al., 2025). However, training with β-glucan also reduces bleomycin-induced lung fibrosis (Y.-Y. Kang et al., 2024). Hence, we aim to optimize relevant intrapulmonary exposures to assess how trained immune responses are modulated by the C3a-C3aR axis, while also clarifying the extent to which central C3-C3aR-mediated training alters AM responses. The results may have clinical implications since trained immunity is linked to vaccine effectiveness (for example, with respect to the Bacillus Calmette-Guérin vaccine) and overall immune resilience (Arts et al., 2018; Kleinnijenhuis et al., 2015). Moreover, humans with C3 deficiency demonstrate impaired responses to vaccination (Kim et al., 2018; Pekkarinen et al., 2015). It is tempting to speculate that local augmentation of C3 and/or C3aR activity – could enhance vaccine effectiveness for hard-to-vaccinate pathogens like *Mycobacterium tuberculosis*.

## MATERIALS AND METHODS

Details including mouse strains, design, models of trained immunity, readouts, and statistical analysis are described as per the ARRIVE guidelines and included in Supplementary Material.

## DATA AVAILABILITY

Sequencing data have been deposited in GEO under the accession code GSE281001 at https://www.ncbi.nlm.nih.gov/geo/query/acc.cgi?acc=GSE281001.

## ACKNOWLEDGEMENTS

We thank Dr. John Atkinson for his feedback, and the Genome Technology Access Center at the McDonnell Genome Institute for help with genomic analysis. The Center is partially supported by NCI Cancer Center Support Grant #P30 CA91842 to the Siteman Cancer Center. This publication is solely the responsibility of the authors.

