## Supplementary Material for "The C3-C3aR axis modulates trained immunity in alveolar macrophages"

**TABLES**

**Table S1. Table of pathways based on Reactome analysis with a list of overlapping genes, when comparing downregulated transcripts between trained C3-deficient and wildtype primary alveolar macrophages.**

**METHODS**

**Experimental model details**

Mice

All animal studies were conducted on an approved IACUC protocol. C57BL/6J (RRID:IMSR_JAX:000664, termed wild-type, or WT) and B6.129S4-*C3^tm1Crr^*/J (RRID:IMSR_JAX:003641, termed C3-deficient, or C3KO) mice were obtained from Jackson Laboratory (Bar Harbor, ME, USA). The mice were maintained as groups in our animal housing facility at the Washington University in St. Louis School of Medicine with a 12 h light-dark cycle, with temperatures between 22-23°C and humidity of 68-72%. C3aR-deficient (termed C3aRKO) mice were provided by Dr. Rick Wetsel at the University of Texas in Houston, were bred in house and have been reported previously by us (Kildsgaard et al., 2000; Sahu et al., 2023).

Experimental Design

The ARRIVE guidelines were followed for reporting of in vivo experiments, and are reported throughout the Methods and Figure Legends. Female and male mice between the ages of 8 and 16 weeks of age were used in these experiments and were age-matched in each experiment. Sample size was determined based on prior studies on acute lung injury, a minimum of 5 mice were used from each genotype (Sahu et al., 2023). Pre-established exclusion criteria for mice included being pregnant or injured (i.e., unanticipated pre-existing wounds from co-housing). Each experiment was repeated at least twice, and both biological and technical replicates were included. No sample or data point from the analysis was omitted.

*In vivo* trained immunity in mice

WT and C3KO mice were administered heat-killed *Pseudomonas aeruginosa* (HKPA, Pa57-15, 1x10^5^ CFU/mouse) (Sahu et al., 2023) or PBS vehicle control intranasally. After 14 days, LPS (*E. coli* O111:B4, Millipore Sigma, cat #: L4391, 10 µg/mouse) or PBS was also intranasally given to these mice. 24 h later, mice were anesthetized with 1.25% tribromoethanol (approx. 125-250 mg/kg) via intraperitoneal injection followed by cervical dislocation for euthanasia. Bronchoalveolar lavage (BAL) was performed by inserting a 20G x 1” Surflo IV catheter into the trachea and instilling approx. 0.5 mL PBS + protease inhibitor (Halt™ Protease and Phosphatase Inhibitor Single-Use Cocktail (100X), Thermo, cat #: 78444) into the lungs. The fluid was then collected, followed by centrifugation at 3,000 rpm for 3 min at 4°C for separating the cell pellet and supernatant, which was then stored at -20°C until later use.

Alveolar macrophage (AM) isolation and maintenance

Ex vivo AM were cultured as per a previously published protocol (Gorki et al., 2022; Zahalka et al., 2022). To obtain AM, BAL was performed at least 4 times per mouse, followed by centrifugation at 3,000 rpm for 3 min at 4°C. The resulting cell pellet was resuspended in AM media (RPMI +10% FBS [Biowest, cat #: S1620], 1% penicillin+streptomycin, 1 µM Rosiglitazone [Millipore Sigma, cat #: R2408], 10 ng/mL mouse TGF-β1 [BioLegend, cat #: 781804], and 30 ng/mL mouse GM-CSF [PeproTech, cat #: 315-03-100UG]), plated on 25 mm round dishes, and incubated at 37°C + 5% CO_2_ for 24 h. Media was then removed, and cells were washed twice with warm PBS to removed unwanted cells and debris. AM media was added again, and cells allowed to grow for at least 7 days. After this, cells were washed again with warm PBS then incubated for 15-20 min with Accutase (ThermoFisher, cat #: MT25058CI) to gently remove them from the plates. They were then centrifuged at 500g for 5 min at 4°C, resuspended in AM media, then placed in T25 flasks and incubated at 37°C + 5% CO_2_. Resulting AMs were passaged every 7 days for a maximum of 25 passages (Gorki et al., 2022).

Induction of trained immune responses in AM in vitro.

AMs were plated at 1x10^5^ cells/well in 96-well flat bottom tissue culture-treated plates and incubated at 37°C +5% CO_2_ for 30 min to 1 h to promote adherence. Each well was then washed with PBS, followed by the addition of “trained” AM media supplemented with HKPA or heat-killed *Candida albicans* (HKCA, 1x10^4^ CFU HKPA or HKCA/well, Invitrogen, cat #: tlrl-hkca) or a matched volume of PBS as a vehicle control for “untrained” wells. Plates were incubated for 24 h at 37°C + 5% CO_2_, washed with PBS, then rested for 6 days in AM media, followed by secondary stimulation with LPS (10 ng/mL, *E. coli* O111:B4, Millipore Sigma, cat #: L4391). After 24 h, supernatants were collected and frozen at -20°C until later use.

Exogenous treatment of AM with C3 and C3a

In order to investigate the effects of C3 uptake or C3a individually on training, either C3 (15 µg/mL, CompTech, cat #: M113) or C3a (10 µg/mL, CompTech, cat #: A118) were added to wells 1 h prior to training of WT, C3KO, or C3aRKO AMs with HKCA. To examine whether C3/C3a affects training via the C3a receptor (C3aR), wells were supplemented with 200 nM of C3aR antagonist SB290157 (Millipore Sigma,cat #: SML1192) 30 min prior to either training with HKCA, or addition of exogenous C3 or C3a followed by training 1 h later.

Quantification of chemokines and cytokines

Untrained and trained BAL and AM supernatants obtained as described above were thawed to room temperature from -20°C. Concentrations of CXCL1, CXCL2, IL-6, and TNFα, or also RAGE and total protein in BAL, were determined via competitive ELISA plates (R&D Systems Inc., Minneapolis, MN, USA) or Milliplex plates (Millipore Sigma, St. Louis, MO, USA) according to manufacturer’s instructions at 1:2 or 1:4 dilution. Sandwich ELISAs were then read on an EPOCH microplate reader via optical density measurement at 450 nm wavelength. Multiplex ELISA plates (Millipore Sigma, St. Louis, MO, USA) were read using a Bio-Rad Luminex 100 multiplex system.

Quantification of C3a-neo

The protocol for measuring C3a-neo in the supernatant was adapted from a previously published protocol measuring it in the serum (Pagano et al., 2009). ELISA plates (96-well flat-bottom; #3855, Thermo Fisher Scientific) were coated with Mouse C3a Capture Antibody (1:250; 100µl per well of PBS; Purified Rat Anti-Mouse C3a, Cat# 558250, BD Pharmingen) overnight at 4°C. After washing three times with a solution of 0.05% Tween 20 in PBS, the plate was blocked with 1% bovine serum albumin (BSA; #A7906, Sigma-Aldrich) at room temperature (RT) for 1 h. Plates were washed again and samples diluted at 1:4 in 1% BSA/PBS solution were added (100 μl per well). Standard curve, made from purified mouse C3a (Purified Mouse C3a Protein, Cat# 558618 BD Pharmingen) was used from 50 to 3.125 ng/ml. Samples and purified protein were incubated at RT for 2 h. The plates were subsequently washed three times and Mouse C3a Detection Antibody (1:1000; 100µl per well in 1% BSA/PBS; Biotin Rat Anti-Mouse C3a, #558251, BD Pharmingen) was diluted in 1% BSA. After another three washes, samples were incubated with Streptavidin HRP-conjugated (100µl per well; 1:200 dilution, #DY998, R&D Systems) for 30 min at RT. After three washes, TMB Color Substrate (#DY999, R&D Systems) was added at 100 μl per well and incubated at RT for 5 min. The reaction was stopped by addition of 1 M sulfuric acid (50 μl per well; #DY994, R&D Systems), and optical density was measured at 450 nm (Epoch Microplate Spectrophotometer, BioTek).

Phagocytosis measurements.

Following experimental treatments, WT and C3KO AMs plated at 1×10⁵ cells/well in 96-well flat-bottom tissue culture-treated plates were washed once with PBS, followed by the addition of pHrodo™ Green E. coli BioParticles (Life Technologies, #P35366) at 25 µg/mL in AM media (100 µL/well). Cells were then incubated at 37°C + 5% CO₂ for 30 min. After this, cells were washed with warm PBS and incubated for 15–20 min with Accutase, then centrifuged at 500 × g for 5 min at 4°C. For viability discrimination, cells were resuspended in Zombie Violet (BioLegend, #423113) at a 1:2,400 dilution in PBS (100 µL/well) and incubated at 4°C in the dark for 15 min. Cells were then washed twice in 1% BSA/PBS and resuspended in 200 µL of 1% BSA/PBS for flow cytometric analysis. Samples were acquired on an Attune Xenith spectral flow cytometer (Thermo Fisher Scientific) operating in conventional mode, using the violet laser (405 nm) for Zombie Violet excitation and the yellow-green laser (561 nm) for pHrodo Green excitation. Phagocytic activity was quantified as the median fluorescence intensity (MFI) of pHrodo Green-positive cells within the live AM gate (Zombie Violet⁻). Data were analyzed using FlowJo v10 (BD Biosciences).

Measurement of Reactive Oxygen Species (ROS) Production.

Following experimental treatments, WT and C3KO AMs plated at 1×10⁵ cells/well in 96-well flat-bottom tissue culture-treated plates were washed once with PBS, followed by the addition of AM media containing CellROX Green (Thermo Fisher Scientific, #C10444) at a final concentration of 500 nM. Cells were incubated for 30 min at 37°C + 5% CO₂ in the dark. After this, cells were washed with warm PBS and incubated for 15–20 min with Accutase, then centrifuged at 500 × g for 5 min at 4°C. For viability discrimination, cells were resuspended in Zombie Violet (BioLegend, #423113) at a 1:2,400 dilution in PBS (100 µL/well) and incubated at 4°C in the dark for 15 min. Cells were then washed twice in 1% BSA/PBS and resuspended in 200 µL of 1% BSA/PBS for flow cytometric analysis. Samples were acquired on Cytek Aurora Spectral flow cytometer (Cytek Bioscience) using a full spectrum unmixing mode. Spectral unmixing was performed using single-color controls and an autofluorescence reference collected from unstained alveolar macrophages. ROS production was quantified as the MFI of CellROX Green within the live AM gate (Zombie Violet⁻). Data were unmixed using SpectroFlo (Cytek Biosciences) and analyzed for downstream gating using FlowJo v10 (BD Biosciences).

Flow Cytometric Measurement of Neutrophils.

BAL fluid was collected as described above and centrifuged at 500 × g for 5 min at 4°C to pellet cells. To remove red blood cells, the cell pellet was resuspended in ACK lysis buffer (150 mM NH₄Cl, 10 mM KHCO₃, 0.1 mM Na₂-EDTA, pH 7.2–7.4) for 2 min at room temperature, then quenched with 1% BSA/PBS and centrifuged at 500 × g for 5 min at 4°C. For viability discrimination, cells were resuspended in Zombie Violet (BioLegend, #423113) at a 1:2,400 dilution in 1% BSA/PBS and incubated at 4°C in the dark for 15 min. Cells were then washed twice in 1% BSA/PBS and stained with CD45-PE/Cyanine7 (BioLegend) and Ly6G-FITC (BioLegend) for 20 min at 4°C in the dark. Cells were washed twice in 1% BSA/PBS and resuspended in 200 µL of 1% BSA/PBS for acquisition. Samples were acquired on an Attune NXT flow cytometer (Thermo Fisher Scientific) operating in conventional mode, using the violet laser (405 nm) for Zombie Violet excitation, the blue laser (488 nm) for FITC excitation, and the yellow-green laser (561 nm) for PE/Cyanine7 excitation. Neutrophils were identified by sequential gating on Zombie Violet⁻ (live) CD45⁺ Ly6G⁺ cells. Data were analyzed using FlowJo v10 (BD Biosciences).

Cell metabolism

To examine a possible mechanistic basis for differences in trained immune responses in WT, C3KO, and C3aRKO AMs, cells were plated at 1x10^5^/well of an XFe24-well cell culture plate (Agilent Technologies, Santa Clara, CA, USA) and trained in AM media with or without C3/C3a, or SB290157 and subsequently washed and rested as described above. After 6 days, cells were restimulated with LPS (10 ng/mL) for 24 h, washed, then left in mitochondrial stress test buffer (Agilent XF DMEM media supplemented with 10 mM glucose, 2 mM glutamine, and 1 mM sodium pyruvate, pH 7.4) at 37°C with no CO_2_ for 1 h. Glycolysis by means of extracellular acidification rate (ECAR) via lactate production in the supernatant was analyzed using an Agilent Seahorse XFe24 analyzer. Three time points were measured each of stable basal glycolysis; after addition of 2.5 µM oligomycin and following the injection of 2 µM FCCP for maximal glycolysis; and was performed according to manufacturer instructions.

Real-time confocal microscopy of live, intact mouse alveoli

*Animals.* Mice were Swiss Webster, purchased from Charles River Laboratories and Taconic Biosciences, 25-40 g, and 6-12 weeks old.

*Solutions*. We purchased Ca^2+^- and Mg^2+^-containing DPBS and Ca^2+^- and Mg^2+^-free PBS from Corning. Isolated mouse lungs were perfused with HEPES-buffered solution of pH 7.4 and osmolality 333 mOsm/L and containing 150 mM Na^+^, 5 mM K^+^, 1 mM Ca^2+^, 1 mM Mg^2+^, 140 mM Cl^-^, 10 mM glucose, 4% dextran (70 kDa; Molecular Probes), and 1% FBS (Gemini Bio-Products). Fluorophores, reagents, and antibodies microinstilled into alveoli were dissolved or suspended in HEPES-buffered vehicle solution containing 150 mM Na^+^, 5 mM K^+^, 1 mM Ca^2+^, 1 mM Mg^2+^, 140 mM Cl^-^, and 10 mM glucose.

*Reagents*. Reagents were freshly constituted for experiments. We purchased calcein red-orange AM (10 μM) and PE-tagged anti-CD11c Ab from from ThermoFisher Scientific. Mouse C3 and C3a were purchased from Complement Technology, incubated with Alexa Fluor-NHS Ester Kit per protocol (Thermo Fisher Scientific) and stored at -80ºC prior to use.

*Preparation of isolated, perfused lungs for microinstillation and imaging*. We anesthetized mice with inhaled isoflurane (4%) and intraperitoneal injections of ketamine (up to 100 mg/kg) and xylazine (up to 5 mg/kg), then gave intracardiac injections of heparin (50 units; Mylan) and exsanguinated the mice by cardiac puncture. We used our established methods (Hook et al., 2018; Tang et al., 2023) to cannulate the trachea, pulmonary artery, and left atrium of the heart, then excise the heart, lungs, and cannulas en bloc. The lungs were positioned to enable micropuncture and imaging of the diaphragmatic surface of the right middle lobe, right caudal lobe, or left lung. Then, we inflated the lungs with room air through the tracheal cannula and perfused the lungs through the pulmonary arterial and left atrial cannulas at 0.5-1.0 mL/min with autologous blood diluted in the lung perfusate solution (see “Solutions”) and warmed to 37ºC. We used in-line pressure transducers (ADInstruments) to maintain constant airway pressure 6 cm H_2_O via a continuous positive airway pressure (CPAP) machine (Philips Respironics) and pulmonary artery and left atrial pressures 10 and 3 cm H_2_O, respectively, via a roller pump (Ismatec). Portions of the lung surface that were not used for micropuncture and imaging were covered with plastic wrap to prevent desiccation.

*Alveolar microinstillation.* We hand-beveled glass micropipettes (Sutter Instruments) to micropuncture single alveoli under bright-field microscopy, as we have done previously (Hook et al., 2018; Tang et al., 2023). Micropunctured alveoli were instilled with fluorophores and reagents in solution, resulting in their spread from the micropunctured alveolus to neighboring alveoli. Microinstillations were performed in 1-3 alveoli bordering each imaging field.

*Live lung imaging and analysis.* Using our established methods previously (Hook et al., 2018; Tang et al., 2023), we viewed alveoli by confocal microscopy (LSM800; Zeiss) with a 20x water immersion objective (NA 1.0; Zeiss) and coverslip. We used bright-field microscopy to randomly select regions of 30-50 alveoli for microinstillation and imaging. All images were acquired as single images using Zen (v.2.6; Zeiss) and recorded as Z-sections. Analyzed images were 4-8 μm below the pleura. Optical thickness was 2 μm, and frame size was 512 x 512 pixels. We established laser, filter, pinhole, and detector settings at the beginning of each imaging experiment to optimize alveolar fluorescence and avoid fluorescence saturation, then maintained the settings for the duration of the experiment. We confirmed absence of bleed-through between fluorescence emission channels. Images were analyzed using ImageJ (NIH; v.2.0.0-rc-69/1.52n). Linear adjustments of brightness and contrast were applied to individual color channels of entire images and equally to all experiment groups. We did not apply downstream processing or averaging.

*Statistics.* Statistics are indicated in figures and legends. We considered statistical significance at *p*<0.05. Data were analyzed and figures were prepared using Microsoft Excel, StatPlus:mac Pro (AnalystSoft, Inc., Build 7.5.0.0/Core v7.6.11), and SigmaPlot (Systat, version 14.5).

*Study approval.* The Institutional Animal Care and Use Committee of the Icahn School of Medicine at Mount Sinai approved the animal procedures.

Human Precision-Cut Lung Slices (PCLS) Preparation

Deidentified human lungs were obtained from OneLegacy. The left lobe was filled with 2% LMP agarose (VWR, 75816-210) kept warm at 40ºC. Once the agarose had congealed, the lobes were excised and placed in ice-cold PBS. Outer pleura lining was removed from the lung lobe, lobes were sliced into ¾-inch cross sections, an 8 mm biopsy punch was used to create samples for sectioning, and airways were avoided due to difficulty with slicing. Punched samples were kept in ice-cold PBS until slicing. Lobes were then embedded for slicing (VF-510-0Z, Precisionary Instruments) at 350 µm in 2% LMP agarose. Slices were placed immediately into cold recovery media and allowed to warm to room temperature before moving to 37ºC incubator. Recovery media was changed after 30 min, 2 h, and overnight incubation. PCLS media was changed each day afterwards.

PCLS Viability

Culture media was collected each day after initial slicing to determine viability through Invitrogen™ CyQUANT™ LDH and G6PD Cytotoxicity Assays (Fisher Scientific, C20301). This data was matched alongside LIVE/DEAD™ Viability/Cytotoxicity Kit, for mammalian cells (ThermoFisher, L3224). Two fresh slices were used each day for this assay. They were first stained for 30 min in 1 mL of 10 µL of Calcein in 10 mL of dPBS; followed by an additional 30 min with 1 µL of Ethidium Homodimer added to each well. Slices were imaged on ECHO Revolve Microscope at 4x. Viability was also confirmed through the presence of ciliary beating in the airways.

Protein Uptake

C3, C3, and C3a (Complement Technology) were first conjugated with Alexa Fluor™ 647 NHS Ester (Succinimidyl Ester) (ThermoFisher, A20006). The final concentration of protein was brought up to be 1 µg/µL. Two PCLS slices were loaded per mouse with 30 µg of C3 for 90 minutes at 37C. An equivalent amount of dye was loaded into wells for each mouse as a control. After incubation, cellular activity was stopped by fixation in 4% paraformaldehyde for 20 min.

Fluorescence Staining

After fixation, slices were washed in PBS and blocked in 5% BSA for 1 h at room temperature. Slices were then washed again in PBS and stained for CD68 (abcam, ab955) in human lungs at 1:100 concentration, followed by washing 3x. Secondary staining was done at 1:200 concentration with Donkey anti-Mouse IgG (H+L) Highly Cross-Adsorbed Secondary Antibody, Alexa Fluor™ 647 (ThermoFisher, A31571), , followed by washing 3x. Counterstaining was done with DAPI at 5 µL in 10 mL of PBS for 10 min. Slices were then kept in 4% PFA overnight before imaging the next day on Zeiss LSM 880 Confocal Microscope. Z-stacks were taken for each slice; boundaries were determined by the presence and absence of DAPI.

Fluorescence Quantification

Quantification was done by a prewritten macro for imageJ to reduce human error in repetitive processing. Each z-stack was converted to a png, each png was measured for green channel (C3/C3a) intensity. ROI was formed by thresholding on CD68 staining and masking (Gaussian Blur sigma=2.5; Thresholding 60-255; Analyze Particles size=400-6000, circularity=0.1-1.00, fill holes). Stacks returning no particles were excluded from analysis. Mean intensity found within ROI was then graphed on Prism.

Single-cell RNA-sequencing (scRNA-seq)

Data from bronchoalveolar lavage (BAL) cells was downloaded from the Gene Expression Omnibus (accession number GSE282132). The dataset included 91,958 cells from 12 samples [DAY2: Saline (3); BCG (3), DAY7: Saline (3); BCG (3)] consisting of BCG and saline-treated individuals at Day 2 and Day 7 after inhalation. The processed H5AD file was imported into R using the zellkonverter package (v1.16.0). Pre-computed dimensionality reduction results, including Harmony-corrected PCA and UMAP embeddings generated by the original authors (Marshall et al., 2025) using harmonypy (v1.2.4), were directly extracted and transferred into a Seurat object (v5.1.0). This ensured that the UMAP visualization matched the published results exactly. Cell type annotations provided in the original dataset were retained for analysis. These included 25 cell populations such as macrophage subtypes (Mo, AcMo, nrMo), T cells, NK cells, dendritic cells, and epithelial cells. Macrophage subpopulations were defined based on canonical marker gene expression as originally annotated in the manuscript: resident alveolar macrophages (Mo) were identified by expression of *MRC1* and *LYZ*; activated macrophages (AcMo) were distinguished by upregulation of *CCL3* and *CCL4*, and non-resident monocytes (nrMo) were identified by expression of *VCAN*, *FCN1*, and *CCL2* (Marshall et al., 2025). To visualize BCG-induced shifts in cell population composition, overlay UMAP plots were generated for each timepoint (Day 2 and Day 7). Complement gene expression was further analyzed across cell populations, with a focus on *C3* and *C3AR1*, to assess the role of the complement system in the early mucosal immune response to BCG infection.

Bulk RNA sequencing

After training C3-deficient and WT alveolar macrophages as described, RNA was extracted using an RNeasy Plus kit (Qiagen, Cat #: 74134). Samples were sent for RNA-Sequencing with polyA selection. Samples were prepared according to library kit manufacturer’s protocol, indexed, pooled, and sequenced on an Illumina NovaSeq 6000. Basecalls and demultiplexing were performed with Illumina’s bcl2fastq software and a custom python demultiplexing program with a maximum of one mismatch in the indexing read. RNA-seq reads were then aligned to the Ensembl release 101 primary assembly with STAR version 2.7.9a (Dobin et al., 2013). Gene counts were derived from the number of uniquely aligned unambiguous reads by Subread:featureCount version 2.0.3 (Liao et al., 2014). Isoform expression of known Ensembl transcripts were quantified with Salmon version 1.5.2 (Patro et al., 2017). Sequencing performance was assessed for the total number of aligned reads, total number of uniquely aligned reads, and features detected. The ribosomal fraction, known junction saturation, and read distribution over known gene models were quantified with RSeQC version 4.0 (Wang et al., 2012).

All gene counts were then imported into the R/Bioconductor package EdgeR (Robinson et al., 2010) and TMM normalization size factors were calculated to adjust for samples for differences in library size. Ribosomal genes and genes not expressed in the smallest group size minus one samples greater than one count-per-million were excluded from further analysis. The TMM size factors and the matrix of counts were then imported into the R/Bioconductor package Limma (Ritchie et al., 2015). Weighted likelihoods based on the observed mean-variance relationship of every gene and sample were then calculated for all samples with the voomWithQualityWeights (Liu et al., 2015) function and were fitted using a Limma generalized linear model with additional unknown latent effects as determined by surrogate variable analysis (SVA) (Leek and Storey, 2007). The performance of all genes was assessed with plots of the residual standard deviation of every gene to their average log-count with a robustly fitted trend line of the residuals. Differential expression analysis was then performed to analyze for differences between conditions and the results were filtered for only those genes with Benjamini-Hochberg false-discovery rate adjusted p-values less than or equal to 0.05.

For each contrast extracted with Limma, global perturbations in known Gene Ontology (GO) terms, MSigDb, and KEGG pathways were detected using the R/Bioconductor package GAGE (Luo et al., 2009) to test for changes in expression of the reported log 2 fold-changes reported by Limma in each term versus the background log 2 fold-changes of all genes found outside the respective term. The R/Bioconductor package heatmap3 (Zhao et al., 2014) was used to display heatmaps across groups of samples for each GO or MSigDb term with a Benjamini-Hochberg false-discovery rate adjusted p-value less than or equal to 0.05. Perturbed KEGG pathways where the observed log 2 fold-changes of genes within the term were significantly perturbed in a single-direction versus background or in any direction compared to other genes within a given term with p-values less than or equal to 0.05 were rendered as annotated KEGG graphs with the R/Bioconductor package Pathview (Luo and Brouwer, 2013).

To find the most critical genes, the Limma voomWithQualityWeights transformed log 2 counts-per-million expression data was then analyzed via weighted gene correlation network analysis with the R/Bioconductor package WGCNA (Langfelder and Horvath, 2008). Briefly, all genes were correlated across each other by Pearson correlations and clustered by expression similarity into unsigned modules using a power threshold empirically determined from the data. An eigengene was then created for each de novo cluster and its expression profile was then correlated across all coefficients of the model matrix. Because these clusters of genes were created by expression profile rather than known functional similarity, the clustered modules were given the names of random colors where grey is the only module that has any pre-existing definition of containing genes that do not cluster well with others. These de-novo clustered genes were then tested for functional enrichment of known GO terms with hypergeometric tests available in the R/Bioconductor package clusterProfiler (Yu et al., 2012). Significant terms with Benjamini-Hochberg adjusted p-values less than 0.05 were then collapsed by similarity into clusterProfiler category network plots to display the most significant terms for each module of hub genes in order to interpolate the function of each significant module. The information for all clustered genes for each module were then combined with their respective statistical significance results from Limma to determine whether or not those features were also found to be significantly differentially expressed. The data was subsequently processed using Partek Flow and pathway analysis was done using EnrichR (Chen et al., 2013; Kuleshov et al., 2016; Xie et al., 2021).

Statistical analyses

Direct comparisons of two isolated groups were analyzed via two-sided unpaired t-test, while multiple two group comparisons were done via t-test with the Holm-Šidák correction to control for the family-wise error rate. Analyses of three or more groups against each other were performed using the one-way analysis of variance with Dunnett’s *post hoc* tests to correct for multiple comparisons. *P* values less than 0.05 were considered statistically significant. Statistical analyses were performed with GraphPad Prism 10.0, and independently with R. Data are shown as individual measurements with mean ± SD, while no outliers have been removed.

Data

The RNASeq data that support the findings have already been deposited under the accession code GSE281001 (<https://www.ncbi.nlm.nih.gov/geo/query/acc.cgi?acc=GSE281001>) and will be made publicly available at the time of publication. The data are available from the corresponding author prior to publication upon reasonable request.
