## Supplementary Figures for "The C3-C3aR axis modulates trained immunity in alveolar macrophages"

### SUPPLEMENTARY FIGURE 1

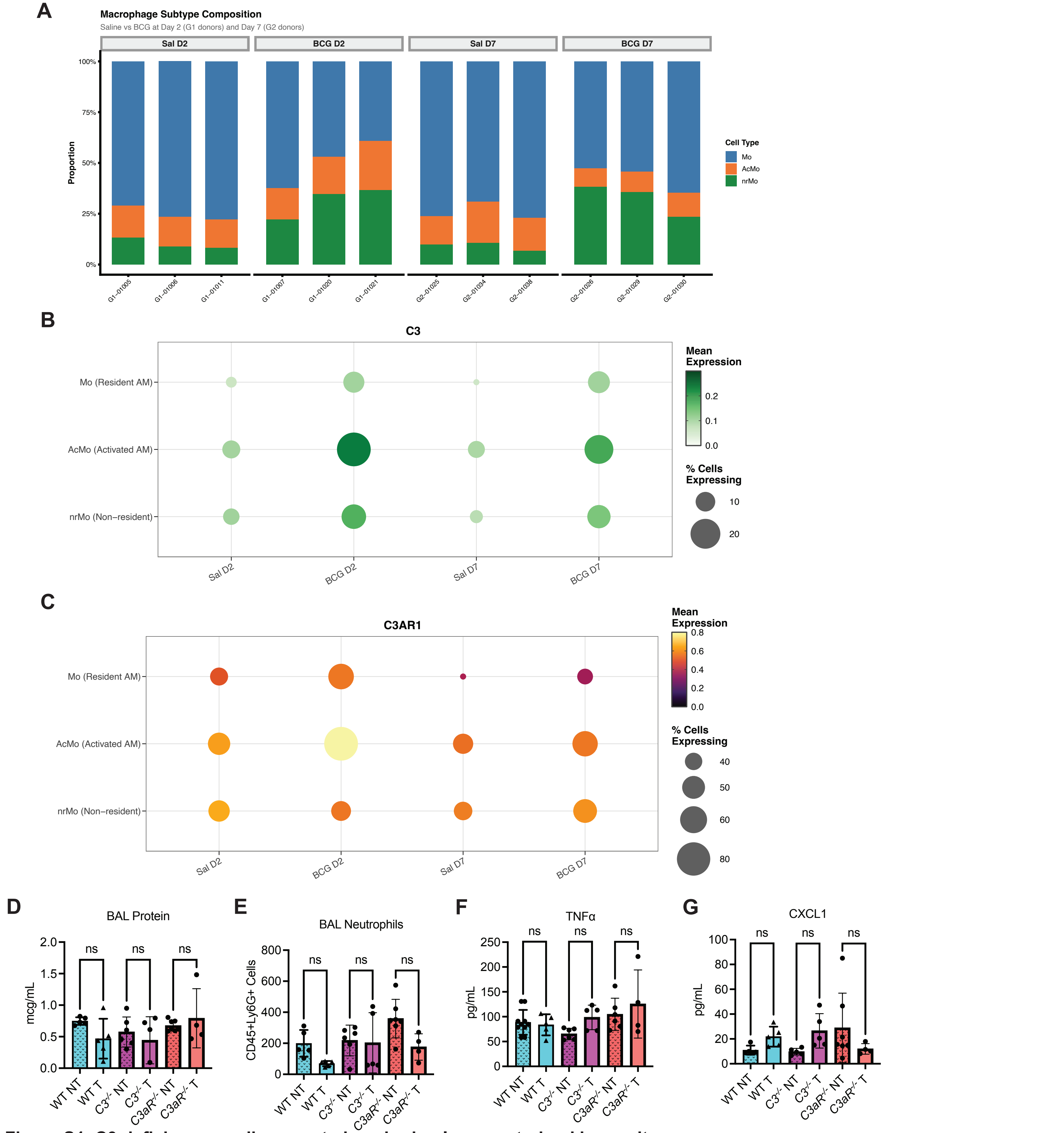


**Figure S1. C3 deficiency predisposes to impaired pulmonary trained immunity.**
(A) Stacked bar charts showing macrophage subtype composition per donor across conditions. Proportions of resident alveolar macrophages (Mo), activated macrophages (AcMo), and non-resident monocyte-derived macrophages (nrMo) are shown for saline (Sal) and BCG-treated donors at Day 2 (G1 donors) and Day 7 (G2 donors). Data from Marshall et al. that included 91,958 cells from 12 samples [DAY2: Saline (3); BCG (3), DAY7: Saline (3); BCG (3)] consisting of BCG and saline-treated individuals at Day 2 and Day 7 after inhalation. (B) Dot plots showing C3 expression across human alveolar macrophage subtypes (Mo, AcMo, nrMo) in saline and BCG-treated donors at Day 2 and Day 7. Dot size reflects the percentage of cells expressing C3; dot color reflects mean normalized expression. Data from Marshall et al. (C) As in (B), for C3AR1. (D-G) Comparison of protein (D), neutrophils (E), TNFα (F) and CXCL1 (G) in the BAL of wildtype (WT), C3-deficient (C3-/-) and C3aR-deficient (C3aR-/-) mice at 14 days after training via the intranasal route with heat-killed Pseudomonas aeruginosa (HKPA). Data were compared with two-sided unpaired t-tests. Each point represents a measurement from one mouse with at least n=4 in each group, mean ± SD shown. *p < 0.05, ns, non-significant.

### SUPPLEMENTARY FIGURE 2

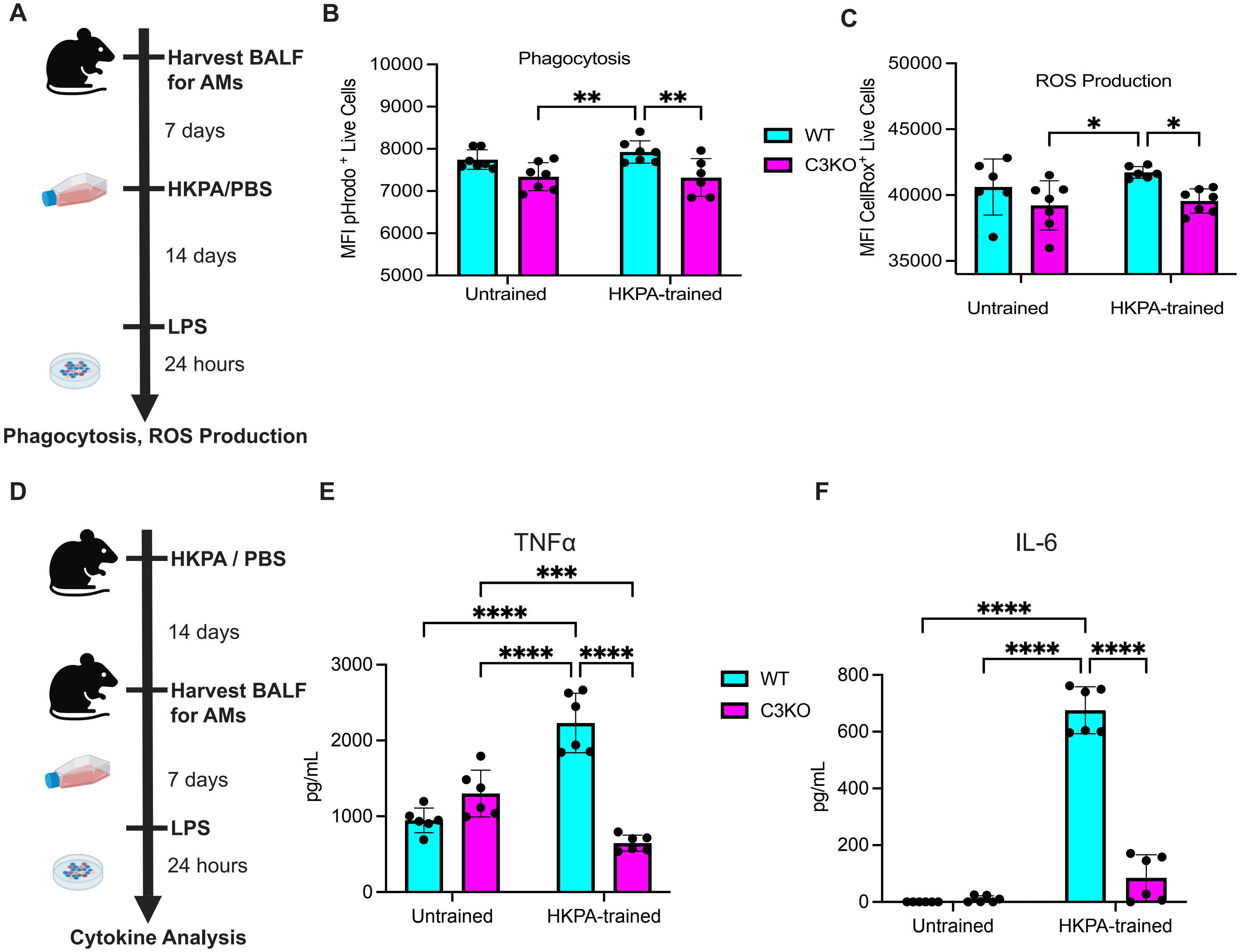


**Figure S2. C3 deficiency results in impaired trained immune responses in LPS-stimulated alveolar macrophages (AMs).**
(A) Schematic representing in vitro training of AMs with HKPA, followed by ex vivo stimulation using LPS and subsequent analysis of the cells. Created with BioRender.
(B) Phagocytic capacity of in vitro HKPA-trained or untrained mouse alveolar macrophages from WT and C3KO mice, measured by mean fluorescence intensity (MFI) of pHrodo-labeled E. coli bioparticles in live cells.
(C) Reactive oxygen species (ROS) production in vitro HKPA-trained or untrained mouse alveolar macrophages from WT and C3KO mice, measured by CellRox mean fluorescence intensity (MFI) in live cells.
(D) Schematic representing in vivo training of mouse lungs with HKPA, followed by ex vivo stimulation of harvested AM using LPS and subsequent cytokine analysis of the supernatants. Created with BioRender. Supernatants were collected at 24 h post-LPS stimulation for cytokine analysis. TNFα (E) and IL-6 (F) concentrations in supernatants from untrained and HKPA-trained WT and C3KO AMs are shown. Data shown as mean ± SD; each point represents one biological replicate. Statistical comparisons by one-way ANOVA with multiple comparisons correction. *$p < 0.05$, **$p < 0.01$, ***$p < 0.001$, ****$p < 0.0001$.

**SUPPLEMENTARY FIGURE 3**

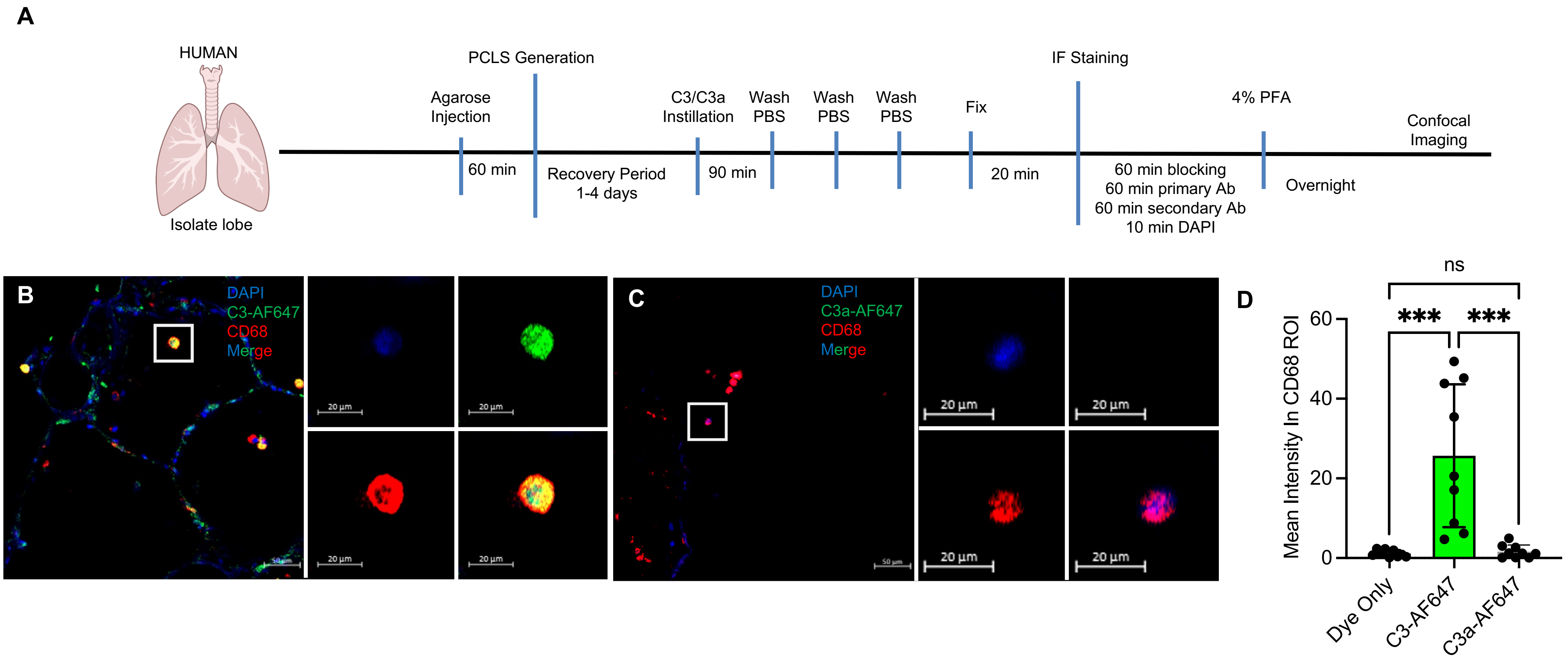


**Figure S3. C3 uptake in human precision-cut lung slices (hPCLS).** (A) Schematic for obtaining human PCLS. Schematic obtained from NIAID Visual & Medical Arts. (10/7/2024). Human Lungs. NIAID NIH BIOART Source. bioart.niaid.nih.gov/bioart/231. (B) Representative image showing C3-AF647 (green) incubated with CD68$^+$ macrophages (red) followed by washing 3x and imaging. Inset shows individual colors. DAPI: blue. (C) Representative image showing C3a-AF647 (green) incubated with CD68$^+$ macrophages (red) followed by washing 3x and imaging. Inset shows individual colors. DAPI: blue. (D) Comparison of individual data points from n=3 hPCLS, 3 technical replicates per donor. Y-axis represents the mean fluorescence within the region of interest (ROI) defined by CD68. Data shown as mean ± SD. Statistical comparisons by one-way ANOVA with multiple comparisons correction. ***$p < 0.001$.
